# Investigating circadian rhythmicity in pain sensitivity using a neural circuit model for spinal cord processing of pain

**DOI:** 10.1101/107375

**Authors:** Jennifer A. Crodelle, Sofia H. Piltz, Victoria Booth, Megan Hastings Hagenauer

**Author notes:** co-first author.

## Abstract

Primary processing of painful stimulation occurs in the dorsal horn of the spinal cord. In this article, we introduce mathematical models of the neural circuitry in the dorsal horn responsible for processing nerve fiber inputs from noxious stimulation of peripheral tissues and generating the resultant pain signal. The differential equation models describe the average firing rates of excitatory and inhibitory interneuron populations, as well as the wide dynamic range (WDR) neurons whose output correlates with the pain signal. The temporal profile of inputs on the different afferent nerve fibers that signal noxious and innocuous stimulation and the excitability properties of the included neuronal populations are constrained by experimental results. We consider models for the spinal cord circuit in isolation and when top-down inputs from higher brain areas that modulate pain processing are included. We validate the models by replicating experimentally observed phenomena of A fiber inhibition of pain and wind-up. We then use the models to investigate mechanisms for the observed phase shift in circadian rhythmicity of pain that occurs with neuropathic pain conditions. Our results suggest that changes in neuropathic pain rhythmicity can occur through dysregulation of inhibition within the dorsal horn circuit.

## 1 The neural processing of pain

The ability for an organism to detect pain is essential for its survival. It is intuitive that the processing of pain must engage a wide-variety of neural circuits ranging from the spinal cord, up through the brainstem, thalamus, and cortex. Though this is true, many of the higher-level cognitive and emotional influences re-converge at the level of the spinal cord, to gate the input of nociceptive information entering the dorsal horn. The dorsal horn serves as a processing center for incoming pain signals, while the midbrain and cortex, as a whole referred to as descending or top-down inhibition [22], serve as a modulator of the pain circuit in the dorsal horn. As a result, there is a tradition of modeling pain processing by focusing exclusively on spinal cord circuitry.

The neural circuit in the dorsal horn receives information about stimulation of peripheral tissues from several types of primary afferent nerve fibers. Nerve fibers called nociceptors detect painful stimuli and are only activated when a stimulus exceeds a specific threshold. These afferents have their cell bodies in the dorsal root ganglia, a cluster of nerve cell bodies located in the back of the spinal cord, and their axons, or afferent fibers, reach to the dorsal horn [3].

There are two major classes of nociceptive fibers: medium diameter, myelinated, fast conducting Aδ fibers that mediate localized, fast pain, and small diameter, un-myelinated, slow conducting C fibers that mediate diffused, slow pain. In addition to the two nociceptive fibers, there are large diameter, myelinated, rapidly conducting A*β* fibers that respond to innocuous, mechanical stimulation [25]. The dorsal horn circuit is composed of many populations of neurons, including excitatory and inhibitory interneurons, and the Wide Dynamic Range (WDR) neurons, or projection neurons. These WDR neurons respond to input from all fibers, and constitutes the majority of the output from the dorsal horn circuit up to the brain.

In this article, we introduce a mathematical model of the pain processing neural circuit in the dorsal horn. We are particularly interested in using the model to investigate mechanisms for circadian and sleep-dependent modulation of pain sensitivity. As reviewed in [9], pain sensitivity exhibits a daily rhythm with a trough in the late afternoon and a peak sometime after midnight for humans. There are several hypotheses for the source of this circadian rhythm, including the sensory afferent fibers, and the top-down inhibition. Since the dorsal root ganglia rhythmically express clock genes responsible for generating circadian rhythmicity of other physiological processes [37], and the dorsal root ganglia contain the cell bodies for a wide variety of afferent neurons, we assume for our model that circadian modulation occurs at the level of primary afferent input to the spinal cord. In the case of more severe or chronic pain, the influences of homeostatic sleep drive on top-down inhibition may be more relevant [37]. We aim to use our mathematical model of the pain circuit in the dorsal horn to form hypotheses on where the modulation might occur and how this placement can affect the firing behavior of the projection neurons.

### 1.1 Previous models of pain processing

There is a long history to understanding how the body perceives pain, including many conflicting theories. Today’s main theory of pain, the gate control theory of pain, was developed in 1965 by Ronald Melzack and Charles Patrick Wall [20]. These researchers revolutionized the understanding of the pain pathway by scrutinizing previous conceptual models of pain processing and developing a model that accounts for the experimental evidence seen thus far. The gate control theory of pain posits that the neural circuitry in the dorsal horn exhibits a gating mechanism that is modulated by activity in the A*β* and C afferent fibers [23]. The nociceptive C fibers facilitate activity in the dorsal horn circuit, whereas the A*β* fibers inhibit activity. When the amount of painful stimuli (activity in C fibers) outweighs the inhibition from the A*β* fibers, the “gate opens” and activates the WDR neurons and thus, the experience of pain. Experimentalists have used this theory to frame their investigations on the types of fibers that project to the spinal cord, as well as the role of different neuron types in the dorsal horn.

Although the gate control theory of pain [20] is a simplification and not a complete representation of the physiological underpinnings of pain [23], it has been a productive starting point for several mathematical and computational models of pain. These models in turn have given insight into the underlying mechanism of pain. The gate control theory was shown to explain several observed phenomena in pain and suggested a possible mechanistic explanation for rhythmic pain (i.e., a sudden change in the input from fast or slow afferent fibers) [6]. Later in [5], the authors considered an excitatory and inhibitory connection from the mid-brain to the inhibitory interneurons and projection neurons, respectively, to be included in the model developed in [6]. This generalization made it possible to take the effect of N-methyl-D-aspartate (NMDA) receptors into account, and therefore, allowed for the resulting model to successfully reproduce the “wind-up” mechanism [5] —that is, an increased level of activity in a neuron that is being repeatedly stimulated [21].

Similarly to [6, 5], more recent models of pain have considered a modeling framework at the level of a single neuron. These biophysically detailed models of pain have been constructed by connecting compartmental models of individual neurons in the dorsal horn according to the circuit architecture proposed by the gate control theory [20]. In these models, the action potential firing of an individual neuron is described by a Hodgkin-Huxley model of membrane current [13] with appropriate membrane dynamics and synaptic strengths based on experimental data. This approach allows for a detailed representation of the geometry and biophysics of each neuron connected to the other neurons via a network whose biophysical behavior and characteristics are then calculated numerically [12]. Such a network model has been previously constructed for the interaction between a deep dorsal horn neuron and Aδ fibers [17], a wide-dynamic range projection neuron [1], and for the dorsal horn circuit between a projection, inhibitory, and excitatory neuron [38]. All these models were validated by showing that they are able to reproduce observed phenomena such as wind-up in the presence of nonzero calcium conductances and NMDA [1, 17, 38]. In addition to wind-up, the model in [38] exhibits also pain inhibition via a response to a stimulus in the A fibers, as has been observed experimentally [36]. On the other end of the modeling spectrum, Arle et al. [2] have constructed a very large-scale, physiologically accurate network model of spinal cord neural circuitry that includes numerous known cell types, their laminar distribution and their modes of connectivity. In addition to simulating pain signaling, the network accounts for the primary motor reflex responses. They applied the model to investigate the mechanisms of pain relief through dorsal column stimulation (DCS), a procedure used to treat neuropathic pain. Their results identify limitations of gate control theory and propose alternate circuitry that more accurately accounts for the effects on nociceptive and neuropathic pain of DCS.

In this work, we take a similar approach to the previous models of pain in terms of the network architecture in the dorsal horn proposed by the gate control theory [21]. However, instead of considering a detailed biophysical model of a single neuron as in [1, 17, 38] or a large-scale network of individual neurons as in [2], we construct equations to describe the population activity of projection, inhibitory, and excitatory neurons in the dorsal horn. As a result, we work with average firing rates of each of the three neuron populations according to the formalism developed in [35]. Therefore, our modeling approach is similar to [6] but we give our model predictions in terms of average firing rates of neuron populations instead of potentials of individual cells.

Thus far, we have not encountered an average firing-rate model for pain in the literature. Our choice of modeling framework and dynamic variables is motivated by our long term aim to integrate a model for pain into an existing model for the sleep-wake cycle constructed in terms of the average firing rate of sleep -and wake-promoting neuron populations. Such a combined sleep-wake-pain model would allow us to test existing hypotheses and ask several biologically motivated questions from the model, for example, on the coupling between sleep, circadian rhythms, and pain sensitivity [7], including the case of chronic pain which is not assessed by the existing biophysical models of pain in [17, 38].

## 2 Mathematical Model

In this section we construct our model for the dorsal horn circuit. We choose a firing-rate model in which we describe the firing rate of the projection, inhibitory interneuron and excitatory interneuron populations in the dorsal horn circuit. The following sections define equations of time evolution for the firing rate of populations, as well as the response functions for each population, arrival times for the afferent fibers, and connectivity between populations.

### 2.1 Equations of time evolution

In our model for pain processing, we focus on the dorsal horn and construct equations for the average firing rate of three interconnected neuron populations in the dorsal horn circuit. We assume that the input to our model is a stimulation of the afferent fibers that has been pre-processed in the dorsal root ganglion. Based on this model input, and on the connections between the neuron populations in the dorsal horn, our model predicts the activity of the projection neurons that then proceeds to the mid-brain (see Figure 1).

**Fig.1.**
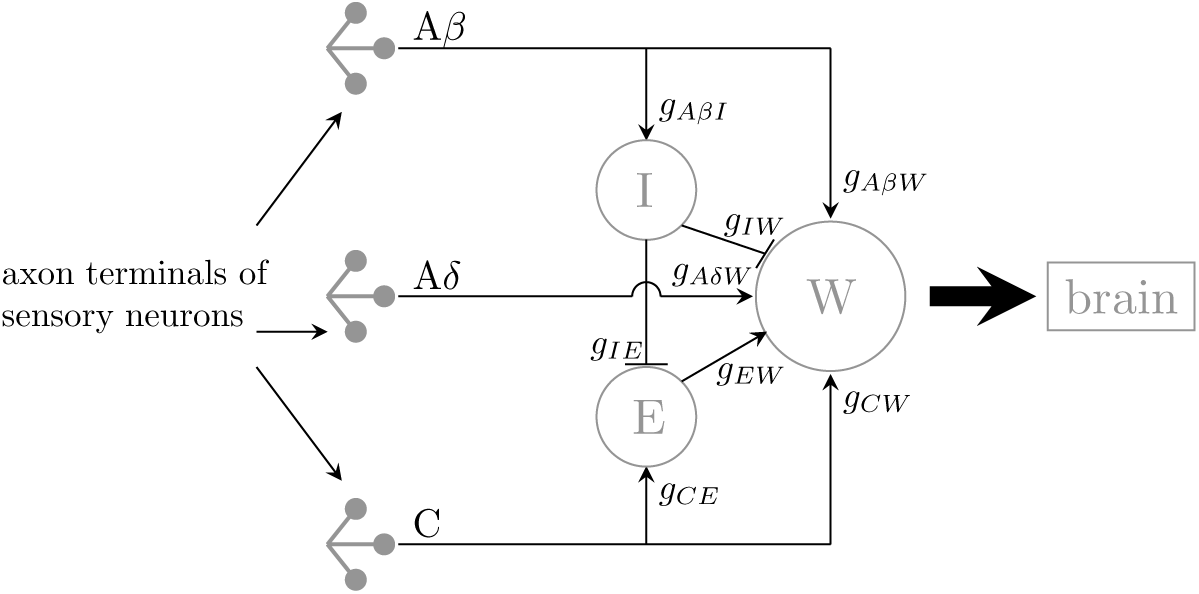
Diagram of our biophysical model for the dorsal horn circuit. For variable names and default parameter values, see Table 1.

In the dorsal horn circuit, the population of the wide-dynamic range (i.e., neurons that respond to both nociceptive and non-nociceptive stimuli) projection neurons (*W*) is connected to the population of inhibitory interneurons (*I*) and excitatory interneurons (*E*) (see Figure 1). According to the formalism of the average firing rate models, we follow [35] and assume that the rate of change of the average firing rate in Hz (i.e., average number of spikes per unit time) of the projection, inhibitory, and excitatory neuron populations, *f*_*W*_, *f*_*I*_, and *f*_*E*_, respectively, is determined by a nonlinear response function (that we define in Section 2.2). These response functions determine the average firing rate of a neuron population based on the external inputs (i.e., stimulations of the afferent fibers pre-processed in the dorsal root ganglion) and the firing rates of the presynaptic neuron populations (see Figure 1). In the absence of input from other neuron populations and afferent fibers, we assume the average firing rate of the neuron population decays exponentially. These assumptions yield the following equations for the average firing rate of each population:

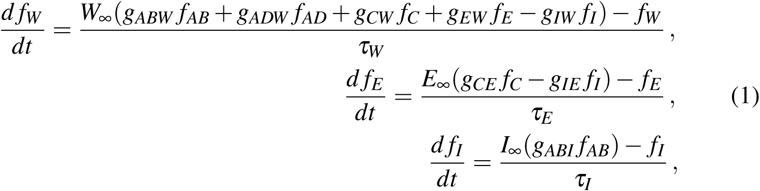

where τ_*W*_, τ_*E*_, and τ_*I*_ are the intrinsic time scales of the projection, excitatory, and inhibitory neuron populations, respectively. Weight *g*_*NM*_ denotes the strength of the effect a change in an external input or presynaptic neuron population *N* has on neuron population *M*. We indicate inhibitory synaptic input with a negative sign and excitatory synaptic input with a positive sign. We define the step functions of the external inputs, *f*_*AB*_, *f*_*AD*_, and *f*_*C*_, and the monotonically increasing firing rate response functions *W*_∞_, *E*_∞_, and *I*_∞_, in the following subsections 2.1.1 and 2.2, respectively. For a summary of all model variables and parameters, see Table 1.

**Table 1.** Summary of our model [in Equation (1)] variables, parameters, and default parameter values.

| Name | Description | Default value |
| --- | --- | --- |
| $g_{ABW}$ | weight of the synaptic connection from $A\beta$ fibers to $W$ | 0.6 |
| $g_{ABI}$ | weight of the synaptic connection from $A\beta$ fibers to $I$ | 0.6 |
| $g_{ADW}$ | weight of the synaptic connection from $A\delta$ fibers to $W$ | 0.3 |
| $g_{CE}$ | weight of the synaptic connection from $C$ fibers to $E$ | 0.6 |
| $g_{CW}$ | weight of the synaptic connection from $C$ fibers to $W$ | 0.4 |
| $g_{EW}$ | weight of the synaptic connection from $E$ to $W$ | 0.4 |
| $g_{IE}$ | weight of the synaptic connection from $I$ to $E$ | 0.05 |
| $g_{IW}$ | weight of the synaptic connection from $I$ to $W$ | 0.15 |
| $\max_{AB}$ | amplitude of the $A\beta$ fiber model input | 2 |
| $\max_{AD}$ | amplitude of the $A\delta$ fiber model input | 0.5 |
| $\max_C$ | amplitude of the $C$ fiber model input | 1.5 |
| $\max_E$ | maximum firing rate of $E$ | 60 Hz |
| $\max_I$ | maximum firing rate of $I$ | 80 Hz |
| $\max_W$ | maximum firing rate of $W$ | 50 Hz |
| $\tau_E$ | intrinsic time scale of $E$ | 20ms |
| $\tau_I$ | intrinsic time scale of $I$ | 20ms |
| $\tau_W$ | intrinsic time scale of $W$ | 1ms |

#### 2.1.1 Model inputs from the dorsal root ganglion

The three different types of afferent fibers not only have different sizes of diameter but they also differ at the level of myelination that provides insulation. As a result, impulses are transmitted at different conductance speeds in the three afferent fibers.

To determine the pattern of nerve input from a painful stimulus to the spinal cord (see Figure 1a in [26]), we simulate the arrival of 1000 nerve impulses. The majority (82%) of these fibers consists of slow C fibers (with an average conduction velocity of 1.25 m/s and a standard deviation of 0.75 m/s), 9% as Aδ fiber fibers (with an average conduction velocity of 0.12 m/s and a standard deviation of 0.083 m/s), and 9% as A*β* fibers (with an average conduction velocity of 0.024 m/s and standard deviation of 0.013 m/s). We assume that the time of initiation of each of the nerve pulses in each of these fibers in the periphery in response to painful stimulation are be roughly equivalent, and that they need to travel 1 meter to reach the spinal cord (e.g., the length of a leg). We choose these proportions and conductance speeds based on the literature [25, 16]. Our simulated data from fibers with different conductance speeds reproduces the observed pattern [26] of a fast response to the A*β*- and Aδ fibers (i.e., first pain) followed by a slow response to the C fibers (i.e., second pain) (see Figure 2).

To generate a simplified model input similar to the simulated input in Figure 2, we use Heaviside step functions to represent how a stimulation (of the afferent A*β* Aδ, and C fibers) and its pre-processing in the dorsal root ganglion is received by the dorsal horn circuit. Thus, the external inputs *f*_*AB*_, *f*_*AD*_, and *f*_*C*_ to the model in (Equation 1 are given by

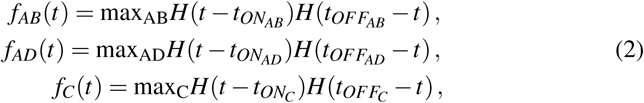

where max_*AB*_, max_*AD*_, and max_*C*_ are the amplitudes of the signals from A*β*, Aδ, and C fibers, respectively, *t*_*ON*_*AB*__, *t*_*ON*_*AD*__, and *t*_*ON*_*C*__ are the time points when an input from A*β*, Aδ, and C fibers, respectively, is received by the dorsal horn circuit, *t*_*OFF*_*AB*__, *t*_*OFF*_*AD*__, and *t*_*OFF*_*C*__ are the time points when an input from A*β*, Aδ fiber, and C fibers, respectively, has decayed and *H(x)* is a Heaviside step function

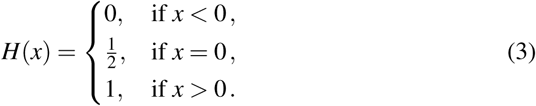

In Figure 3, we show an example input that mimics the combined signal from the three afferent fibers and that we use as an input to our model in Equation (1).

**Fig. 2.**
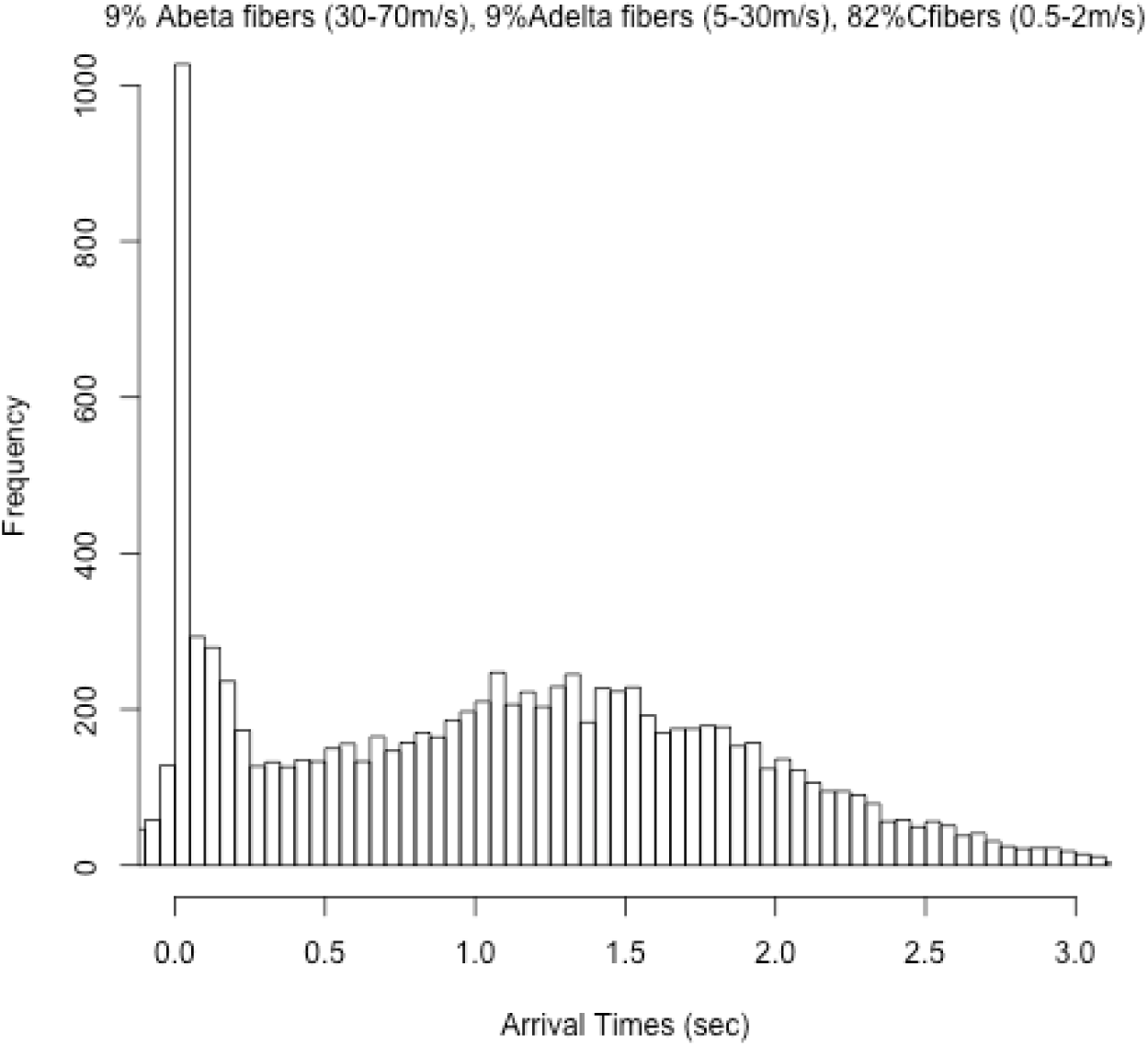
Simulated post-stimulus histogram from three afferent fibers with different conductance speeds reproduces a pattern observed in the action potential of projection neurons [26] where fast pain response is composed of a response to stimuli in the A*β*-and Aδ fibers followed by slow pain response to stimuli in the C fibers.

### 2.2 Firing rate response functions

In our modeling framework, we assume a sigmoidal shape for the monotonically increasing firing rate response functions *W*_∞_, *E*_∞_, and *I*_∞_, and use hyperbolic tangent functions to represent them

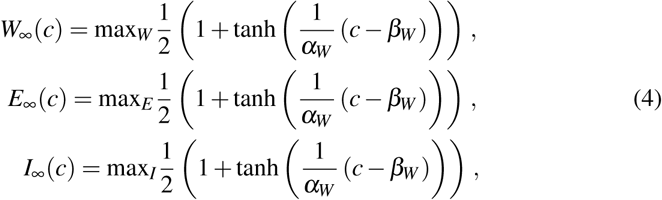

**Fig. 3.**
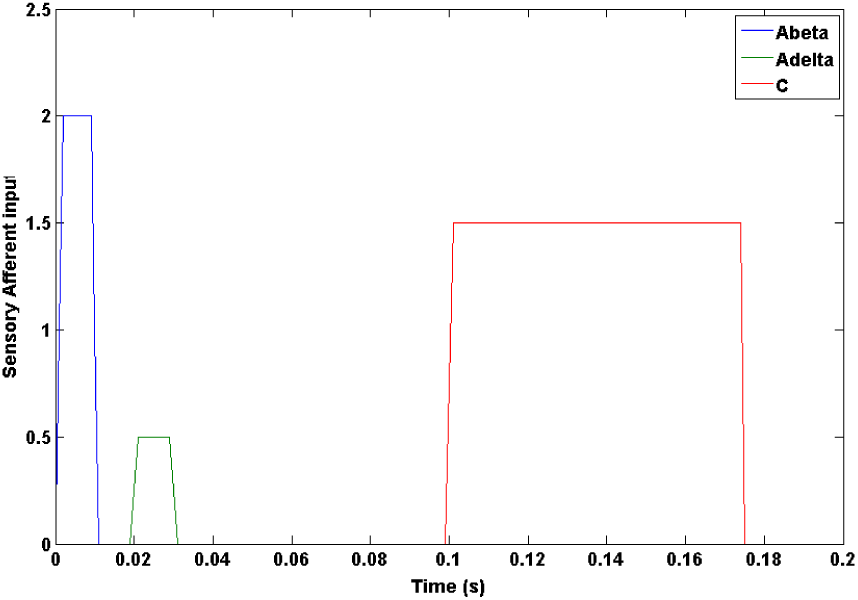
Simulated model input to the dorsal horn circuit from the afferent fibers after pre-processing in the dorsal root ganglion.

where max_*W*_, max_*E*_, and max_*I*_ are the maximum firing rates of the projection, excitatory, and inhibitory neuron population, respectively. In Equation (4), the shape of the tanh-functions is determined by the input *c* at which the average firing rate of the projection, excitatory, and inhibitory neuron population reaches half of its maximum value, *c* = *β*_*W*_, *c* = *β*_*E*_, and *c* = *β*_*I*_, respectively. The slope of the transition from non-firing to firing in the projection, excitatory, and inhibitory neuron population is given by 1/α_*W*_, 1/α_*E*_, and 1/α_*I*_, respectively. See Table 1 for default parameter values. We choose the parameter values for the tanh-functions in such a way that the input-output curve of the projection, excitatory, and inhibitory neuron population agrees qualitatively with experimental observations (see Figure 4). Hence, we assume the inhibitory interneuron population has a nonzero resting firing rate, as has been reported in [3, 22], and a higher maximum firing rate than that of the projection and excitatory interneuron populations, as has been assumed in a biophysically detailed model of the dorsal horn circuit [38]. In our model assumptions for the response functions, we mimic the model predictions of [38] that agree with data from experimental observations in [18, 27]. Thus, we assume that for a small input, the excitatory interneuron population has a small average firing rate that, however, reaches a higher maximum than that of the projection neuron population for a large input (see Figure 4).

**Fig. 4.**
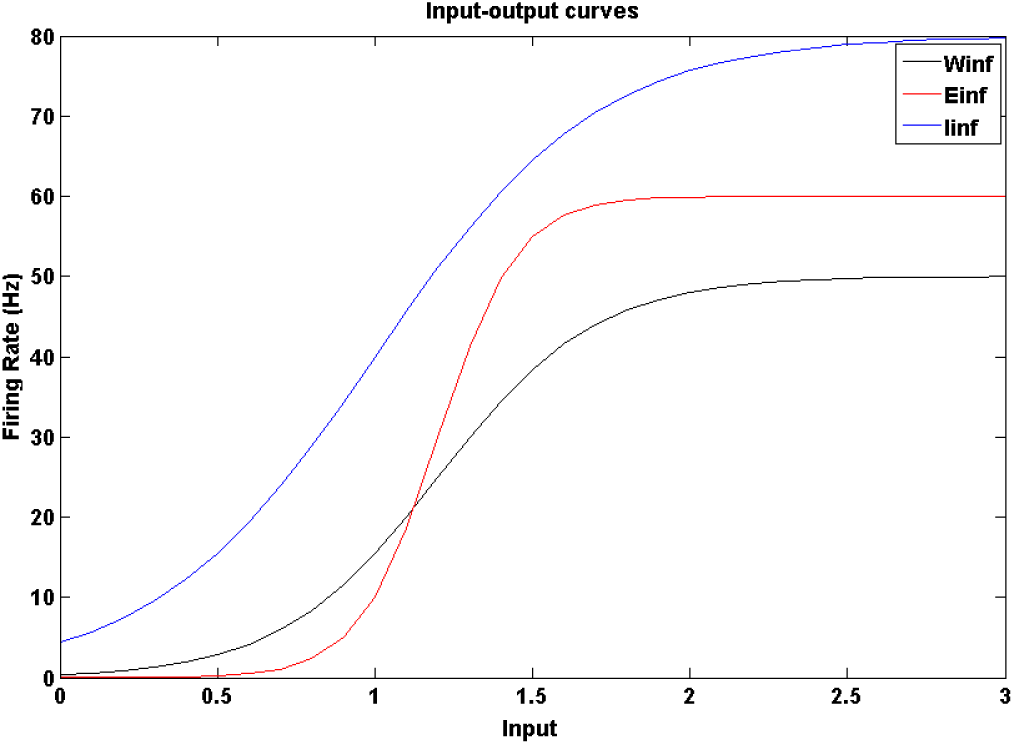
Response functions of the projection (black), excitatory (red), and inhibitory (blue) neuron populations for different constant inputs (on the x-axis). For parameter values, see Table 1.

## 3 Model validation

In this section, we set out to show that our model reproduces various experimental observations such as pain inhibition, wind-up and neuropathic phase changes.

First, we show that our model reproduces the average firing-rate pattern of the populations of neurons in the dorsal horn when the three afferent fibers differ in their conductance speeds. That is, as a response to the input from the afferent fibers as shown in Figure 2, the average firing rates of the projection and interneuron populations [which are connected to each other as shown in Figure 1 and whose dynamics are modeled as in Equation (1)] are qualitatively similar to the simulated histogram in Figure 2 and also seen experimentally [e.g., see Figure 1a in [26] (see Figure 5)]. In addition, the model captures the expected tonic firing rate in the inhibitory neuron population [3, 22], as well as captures the low firing rate of the excitatory neurons [19, 28] (see Figure 4). We use the model output shown in Figure 5 as our point of comparison when choosing “default” values for the weights *g*_*NN*_ (see Table 1) representing the strength of the connections between the neuron populations as shown in Figure 1.

**Fig.5.**
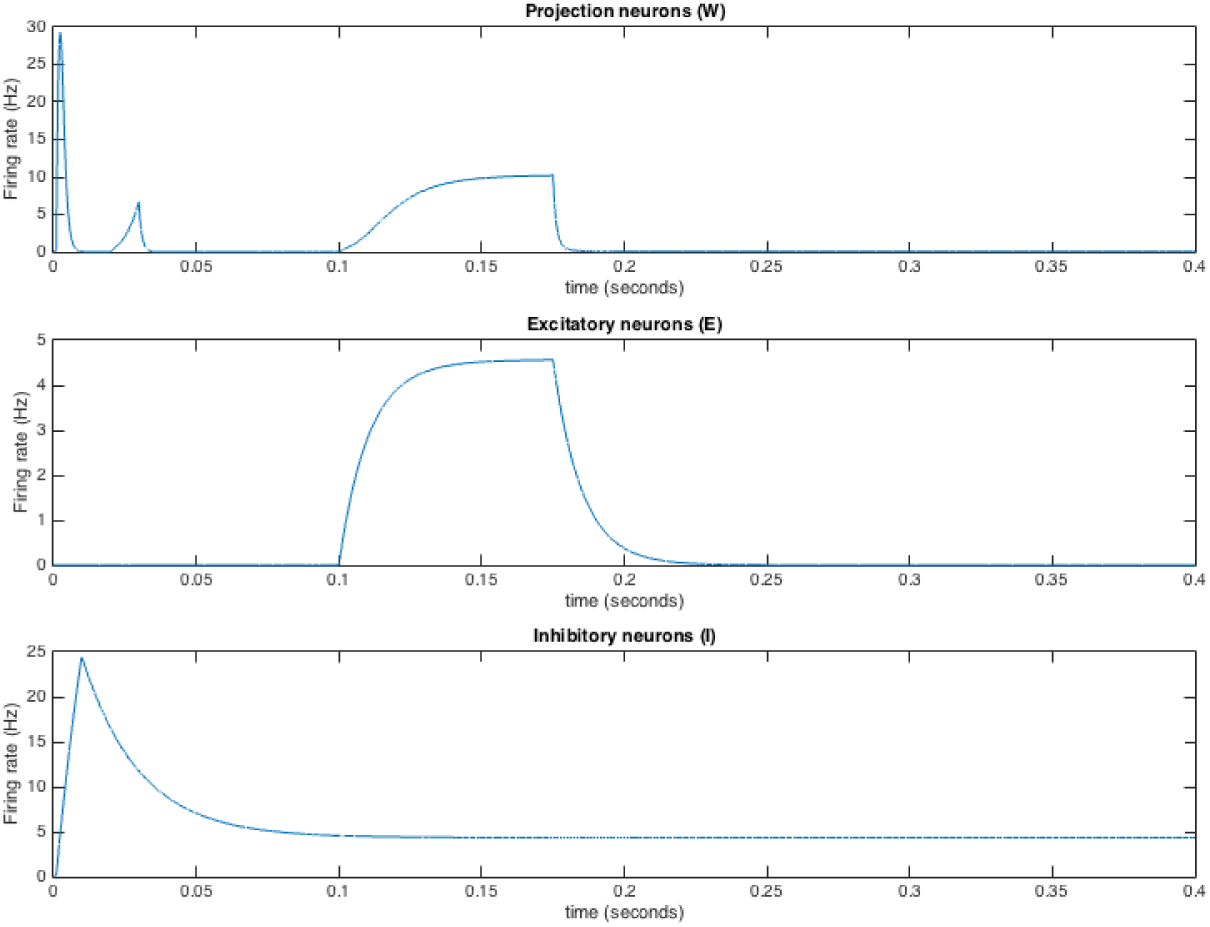
Firing rates for each population in response to the input from afferent fibers as described in Figure 3

### 3.1 Pain inhibition

It has been experimentally observed that stimulation of A fiber afferents can lead to inhibition in some wide-range projection neurons that typically follows from a stimulation of the C fiber afferents [38]. This is related to the idea that when you stub your toe, you immediately apply pressure on the toe and feel some lessening of pain. To capture this phenomenon in our model, we stimulate all three fibers (stubbing of the toe) and then deliver a pulse to the A*β* fiber a short time thereafter (pressure applied to toe), shown in Figure 6 by the red arrows. The arrival time of the second pulse to the A*β* fiber is increased by 10ms for each simulation, and the response in the projection neurons is shown in blue. As can be seen in Figure 6, the timing of the second pulse gets closer to the arrival of the C fiber stimulation, and there is a brief period of excitation followed by a longer period of inhibition, as seen in experiments [36]. Thus, our model successfully captures this delayed inhibition phenomenon.

**Fig. 6.**
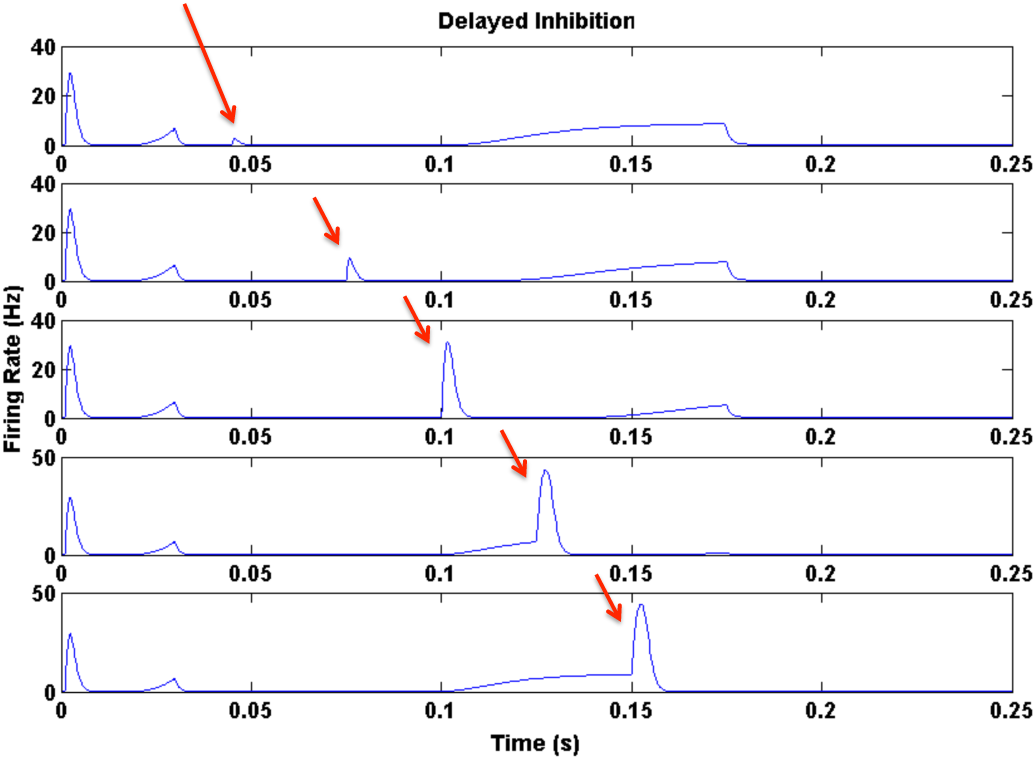
Pain inhibition phenomenon captured in the model. Response of the projection neuron population to the initial fiber pulse stimulation (at *t* = 0) and the second pulse stimulation only to A*β* fibers (red arrow) for increasing in time between fiber stimulations.

### 3.2 Wind-up

We aim to further validate our dorsal horn circuit model (1) by showing that it reproduces wind-up —that is, increased (and frequency-dependent) excitability of the neurons in the spinal cord because of repetitive stimulation of afferent C fibers [21]. Wind-up serves as an important tool for studying the role the spinal cord plays in sensing of pain, and it has been often used as an example phenomenon to validate single neuron models of the dorsal horn (see [17, 1, 38], for example). However, both the physiological meaning and the generation of wind-up remain unclear (see [11] for a review).

There are several possible molecular mechanisms proposed for the generation of wind-up (see Figure 6 in [11]). Earlier work on single neuron models suggests that wind-up is generated by a combination of long-lasting responses to NMDA and calcium currents providing for cumulative depolarization [1]. Indeed, calcium conductances and NMDA receptors of the projection/deep dorsal horn neurons are included in all previous models of the dorsal horn [38, 17, 1]. In contrast to the model in [1], wind-up can also be reproduced in the absence of synapses that express gamma-Aminobutyric (GABA) from C fibers to the projection neuron [38]. The study done in [17] emphasizes the effect (direct or via influencing the dependence of the deep dorsal horn neurons on their intrinsic calcium currents) NMDA and inhibitory conductances have on the extent of wind-up in the deep dorsal horn neurons [17].

Experimental data on superficial and deep dorsal horn suggest that wind-up is exhibited more by the deep than by the superficial dorsal horn neurons [30]. However, wind-up in the potential of the C fibers is observed in the superficial but not in the deep dorsal horn [29]. Similarly to [1], we investigate whether wind-up of the wide-dynamic range projection neurons in the dorsal horn circuit can be explained by an increase in the C fiber response before the C-input reaches the dorsal horn circuit. Thus, we assume wind-up occurs “upstream” from the dorsal horn circuit described by our model in (1), and represent it as an increase in the duration, and as a decrease in the arrival time, of the C fiber model input to the dorsal horn circuit.

Increase in C fiber synaptic efficacy has been proposed as a possible generation mechanism for wind-up in the literature [29] and suggested as one of the molecular mechanisms underlying wind-up (see Fig. 6 in [11]). Similarly to [1], our model predicts an increase in the activity of the projection neurons for an increase in the width of the step input from C fibers (see Figure 7, left). Furthermore, as in [1], our model also predicts that wind-up in the excitatory interneurons (as a response to the change in the C fiber model input) is similar to that seen in the projection neurons (see Figure 7, right). However, such behavior of the excitatory interneurons is not well-supported by experiments where wind-up is mostly observed in the projection neurons [30]. Because wind-up in the excitatory interneurons had not been reported by 2010, C fiber presynaptic facilitation was discarded as a possible mechanistic explanation for wind-up in the modeling work done by [1]. Nonetheless, the authors note that there is a possibility for underestimating the extent of wind-up in interneurons because they are smaller in size than projection neurons, and therefore, more difficult to sample for electrophysiology experiments than projection neurons [1].

We note here that the proposed mechanism we simulate in Figure 7 involves changing the profile of the model input (in Figure 3) which leads to an obvious change in the model output. We discuss implementations of dynamic wind-up mechanisms in our conclusions in Section 5.

**Fig. 7.**
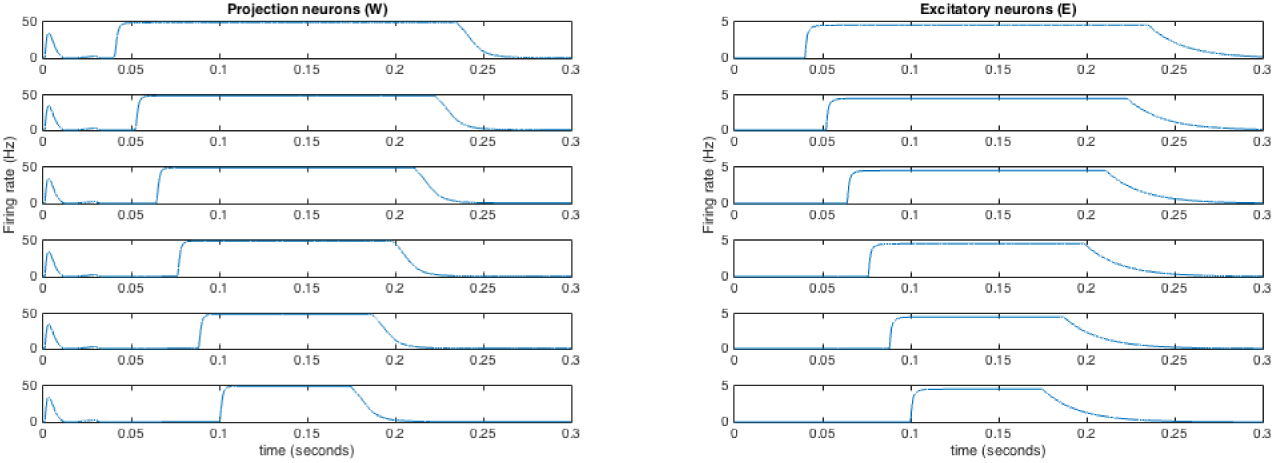
(Left) Projection and (right) excitatory interneuron activity predicted by the model in (1) and otherwise as shown in Table 1] when C fibers are stimulated repeatedly. We assume that repeated stimulation of C fibers is experienced in the neurons upstream from the deep dorsal horn, and thus, seen as an increase in the duration of the C fiber model input (see Figure 3) to the dorsal horn circuit (1). In other words, the model input from C fibers arrives to the dorsal horn circuit at time *t*_*ON*_*C*__ = 0.01s + *n*0.012*s*, where *n* = 0,1,2,3,4,5 and *n* increasing from bottom to top panel.

### 3.3 Neuropathy

Following model creation, we next set out to determine whether changes in the balance of excitation and inhibition within spinal cord pain circuitry could explain changes in pain processing under pathological conditions. We were particularly interested in the case of inflammation and neuropathic pain, in which non-noxious mechanical stimuli become painful following peripheral nerve injury. Both nerve injury and inflammation can cause a deregulation of chloride ion transporters in the dorsal horn. Maintaining a low intracellular chloride concentration is important for the functioning of inhibitory neurotransmission. Under typical conditions, the neurotransmitter GABA produces an inhibitory post-synaptic response by binding to the GABA_*A*_ receptor, which allows negatively-charged chloride ions to flow into the post-synaptic neuron, thus producing hyperpolarization. Under neuropathic or inflammatory conditions, intracellular chloride concentrations may stay semi-permanently elevated, allowing chloride ions to flow out of the cell in response to GABA_*A*_ receptor activity, producing excitatory rather than inhibitory effects. Several authors have hypothesized that this deregulation of spinal pain inhibition could explain the development of pain sensation in response to non-noxious stimuli under neuropathic conditions [34, 10].

Neuropathic conditions are characterized by an 8-12 hour shift in the phasing of daily rhythms in pain sensitivity [37, 31, 15, 8]. As an application of the model, we investigate whether a large phase shift could be produced by a combination of deregulated neural inhibition, and differentially-phased rhythmic afferent input from *Aβ* and C fibers [24], see [9].

Several inflammatory pain conditions, such as osteoarthritis and rheumatoid arthritis, have been shown to exhibit circadian rhythm in pain, with the peak of pain intensity being felt during the night, see Figure 1 in [8]. Neuropathic pain occurs from various conditions involving the brain, spinal cord and nerves. It is distinguished from inflammatory conditions, like arthritis, in that it often appears in body parts that are otherwise normal under inspection and imaging, and is also characterized by pain being evoked by a light touch. Experiments on pain in neuropathic patients suggest that neuropathic pain has a circadian rhythm as well, having its peak in the afternoon, see Figure 2 in [8]. An afternoon peak in pain sensitivity is opposite of the daily rhythm in pain sensitivity under normal conditions [9]. We use our model to further investigate this phenomenon and propose that a possible mechanism for this shift in rhythm is due to the interaction between the *Aβ* and C fibers.

It has been seen experimentally that the A fiber activity can have an inhibitory influence on C fibers, and that under neuropathic conditions, this inhibition can turn to excitation [33, 10]. Using both of these experimentally observed results, as well as the idea that the circadian rhythm comes into the dorsal horn at the level of the fiber inputs we show that we can get a change in phase of the firing rate of the projection neurons with a change from inhibition (normal conditions) to excitation (neuropathic conditions) in the influence from the A fibers to the C fibers. In order to test our hypothesis that under neuropathy, response to acute phasic pain peaks in the late afternoon when, under normal conditions, pain sensation reaches its minimum value, we introduce two principal modifications to our model in Equation (1). First, we impose a circadian rhythm on the maximum amplitude of the model inputs from A*β* and C fibers (i.e., on parameters max_AB_ and max_C_, respectively). Second, we assume an amplitude modulation of the C fibers by the A*β* fibers (see Figure 8).

Our motivation for the second assumption comes from experimental data suggesting that A fibers can decrease the activity of C nociceptors [33]. To represent such an inhibitory effect of A fibers, we model the amplitude modulation between the A*β* and C fibers with a weight *g*_*AβC*_, which under normal conditions is inhibitory and *g*_*AβC*_ < 0, whereas under neuropathy, the inhibitory interneuron population through A*β* fibers has an excitatory effect on the C fibers and *g*_*AβC*_ > 0. By simulating our modified model, we investigate whether the activity of the projection neurons follows the circadian rhythm in C fibers under normal conditions and that of the A*β* fibers under neuropathic conditions.

Earlier work suggests circadian rhythmicity in both the touch and pain sensitivity (see Figures 1 and 2 in [24]). Namely, the pain sensitivity is at its lowest in the early afternoon and at its highest in night, while the highest sensitivity for tactile discrimination is reached in the late afternoon and the lowest in the late morning [24]. These experimental observations motivate us to introduce a circadian rhythm to the model input from *Aβ* fibers that is in antiphase with the circadian rhythm of the C fiber-model inputs, while keeping the arrival times from the three afferent fibers at their default values (see Figure 3 and Table 1). Thus, in our modified model for neuropathy, 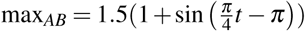

and
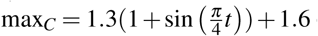
 (see blue and red curves, respectively, in Figure 9). In addition, because of the synaptic connection from the inhibitory interneuron populations to C fibers, we compute the effective maximum height of the C fiber model input as

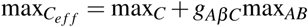

where *g*_*AβC*_ = −0.5 under normal conditions, and *g*_*AβC*_ = 1.5 under neuropathic conditions. In order to assess the extent of experienced pain, we compute the integral of the firing rate of the projection neurons to a stimulus in C fibers (i.e., painful stimuli) using the trapezoidal method from *t* > 0.07*s* onwards (i.e., the C-response of *W*, see model output curve in Figure 5). Indeed, our model simulations suggest that inhibition turned excitation at the level of the fibers is a possible mechanistic explanation for the flip in phase of pain sensitivity seen under neuropathic conditions. Our model shows that under normal conditions, the pain sensitivity rhythm follows the circadian rhythm of the C fibers (see top and bottom panels in Figure 9) but mimics the rhythm in the A*β* fibers under neuropathic conditions (see middle and bottom panels in Figure 9).

**Fig. 8.**
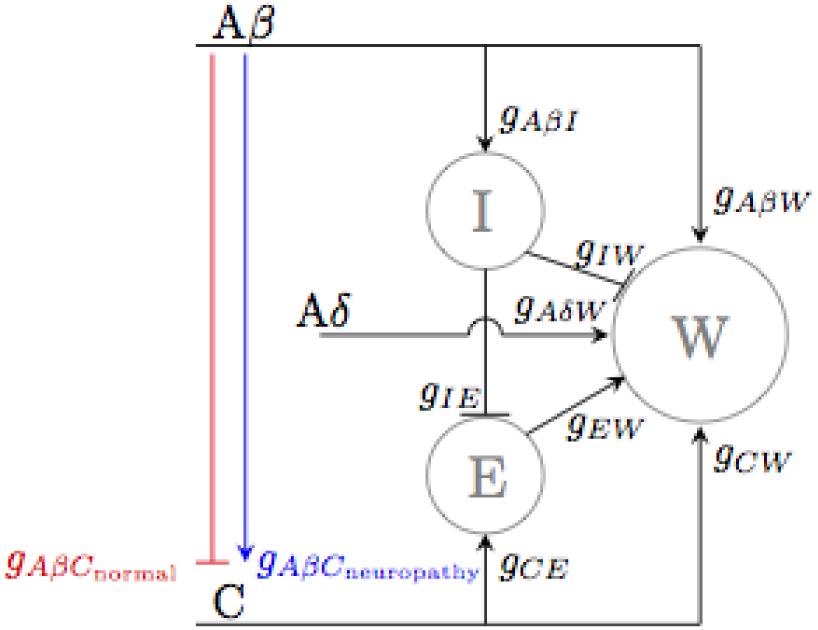
Diagram of our biophysical model for the dorsal horn circuit under (blue) neuropathic and (red) normal conditions.

We note here that under neuropathic conditions, the firing rate of the projection neurons is at, or near, its maximum value throughout the day. While patients with neuropathic pain experience an increased level of pain throughout the day than those without, it is unrealistic for the pain to be at, or near, its maximum all day. In the next section, we propose an amended model in which we include top-down inhibition from the mid-brain where this is not the case.

**Fig. 9.**
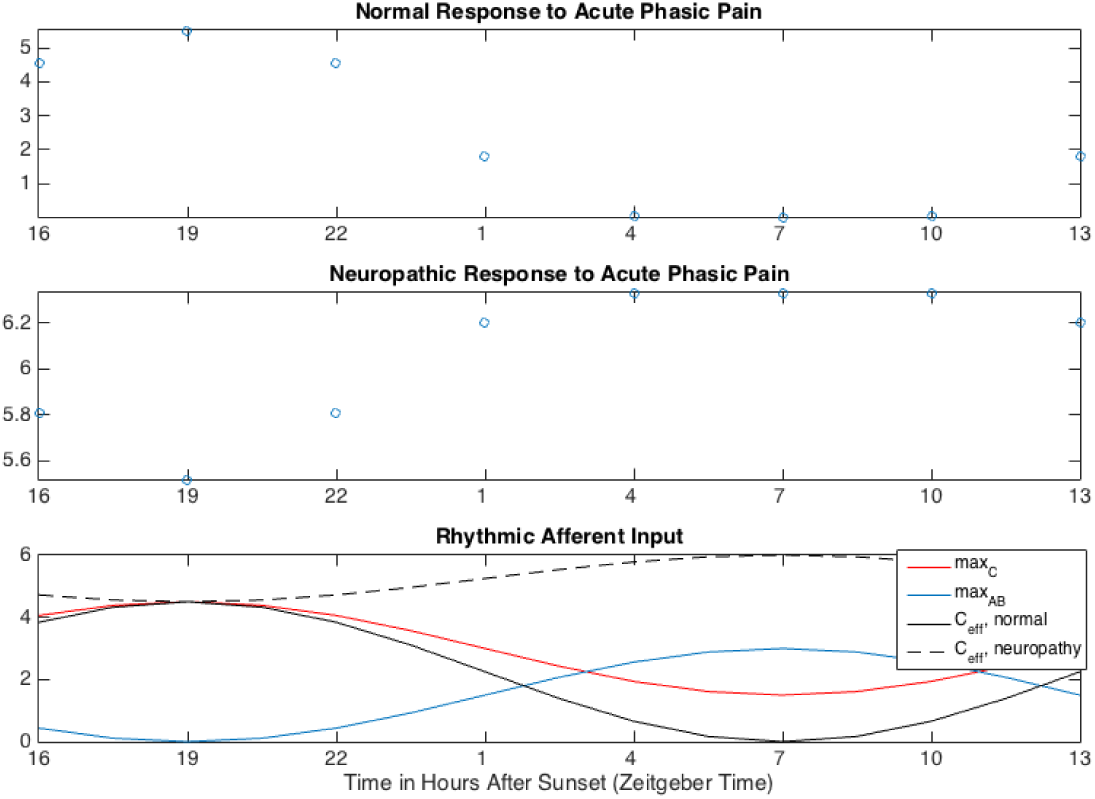
(Circles) Integrated projection neuron activity predicted by the model in (1) under (top) normal and (middle) neuropathic conditions, and (bottom) the different circadian rhythms of the height of the afferent fiber-model inputs. Under normal conditions, the interneuron population *I* decreases the height of the model input from C fibers (see black solid curve in the bottom panel), whereas under neuropathic conditions the connection between *I* and C fiber model input is excitatory. As a result, the effective height of the C fiber model input (black dashed curve) is higher than its baseline value (red curve). We calculate the activity of the projection neuron population as the area under the C-response. Thus, for each zeitgeber time point (with the corresponding maximum heights of the model inputs from A*β* and C fibers shown in the bottom panel), we simulate the neuropathy model for 0.4s and determine the integral under the projection neuron response for *t*= [0.007*s*; 0.4*s*].

## 4 Model with descending control from the mid-brain

### 4.1 Introduction

In its current form, our biophysical model of the dorsal horn pain circuit includes response functions for the three neuron populations that mimic empirical observations. Importantly, our model reproduces the phenomenon of pain inhibition in which a brief mechanical stimulus applied after a painful stimulus can decrease the activity of the projection neurons, and thereby, decrease the sensation of pain. Our model also captures the phase shift in pain intensity for neuropathic pain, however, the amplitude of the pain intensity under neuropathic conditions is very small, and the firing rate of the projections neurons is basically at its maximum throughout the day. In an attempt to both make the model more realistic, as well as explain the neuropathic phase flip, we introduce an amended model in which we consider communication from the dorsal horn to the mid-brain (see Figure 10). Influence from the mid-brain to the dorsal horn plays an important role in modulating inhibition within the pain circuit of the dorsal horn [5, 37, 22]. There are several descending pathways from the brain down to the spinal cord that could affect the afferent fibers, the inhibitory interneurons and the projection neurons, see Figure 2 in [22]. We choose to model the inhibitory descending pathway, as done in [5]. The motivation for this added mechanism is to enable the projection neurons to exhibit a more realistic phase flip in pain sensitivity throughout the day under neuropathic conditions. We also aim to include a mechanism for the effect of the homeostatic sleep drive on this top-down inhibition. As shown in [14] and [32], the build up of the homeostatic sleep drive is reflected in the daily rhythm of the pro-inflammatory cytokines, whose increased levels are associated with an increased firing rate of the WDR projection neurons. We model this by assuming that the connection from the mid-brain to the dorsal horn circuit is a function of the time spent awake, or the build-up of the homeostatic sleep drive. We verify this amended model by showing that it can reproduce the same phenomena as the earlier model, as well as show that this amended model can better capture the observed change in phase of pain sensitivity rhythm for neuropathic patients.

**Fig. 10.**
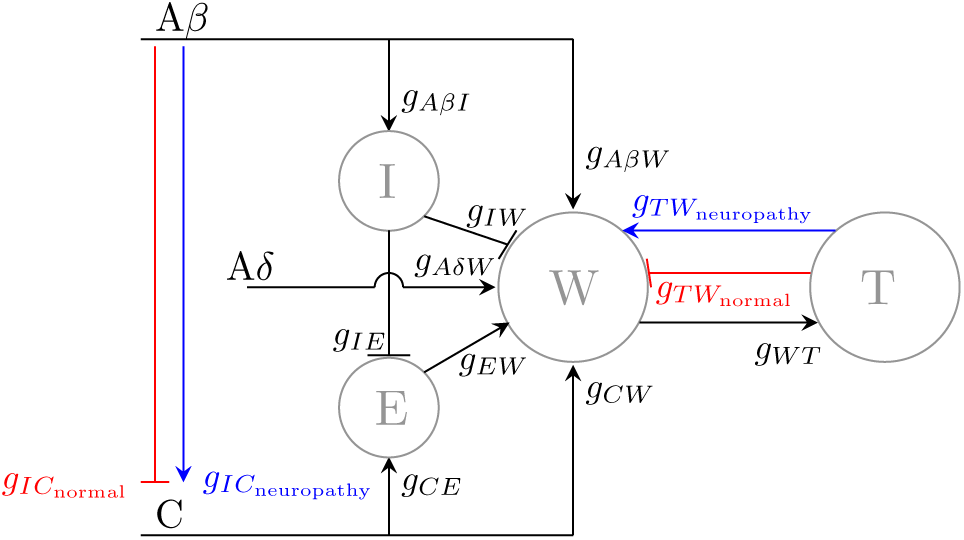
Diagram of our biophysical model for the dorsal horn circuit including connections to and from the mid-brain under (blue) neuropathic and (red) normal conditions.

### 4.2 Amendments to model

In our modified biophysical model for pain, we add a connection between the projection neurons and the mid-brain as shown in Figure 10 and a dimension to the mathematical model in Equation (1). Thus, the dynamics for the average firing rate of neuron populations *I* and *E* remain as they are in Equation (1), and the equations for the projection neurons *W* and neuron population in the mid-brain (*T*) become as follows:

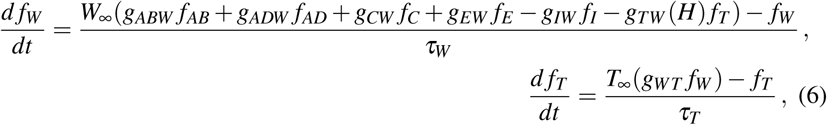

where τ_*T*_ is the intrinsic time scale of the population *T* and weight *g*_*WT*_ denotes the strength of the effect a change in the *W* population has on the neuron population *T*. To investigate the coupling between neuropathic pain and sleep deprivation, we allow, *g*_*WT*_, the strength of the effect of a change in the *T* population on the *W* population, to depend on the homeostatic sleep drive *H*. Hence, we write the weight of the connection from *T* to *W* as *g*_*TW*_ (*H*). As in the case of the other neuron populations in Section 2.2, we assume a monotonically increasing firing rate response function (with respect to input *c*) for the mid-brain population *T*:

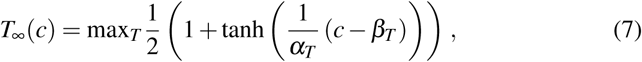

where max_*T*_ is the maximum firing rate of the mid-brain population, *c* = *β*_*T*_ is the input at which the average firing rate of the mid-brain population reaches half of its maximum value, and 1=α_*T*_ determines the slope of the transition from non-firing to firing in mid-brain population.

### 4.3 Model Validation

In regard to parameter values, we introduce a lag in the connection from the midbrain to the projection neurons, and choose τ_*T*_ = 0.05 ms, which is larger than the intrinsic time scales of the other populations (see Table 1). We do this as a result of the assumption that the signal must travel much further to interact with the midbrain than it would to other populations within the dorsal horn circuit. In addition, we assume that the maximum amplitude and the slope of the response function of *T* are smaller than those of the other neuron populations of the model and we pick (max_*T*_; α_*T*_; *β*_*T*_) = (30,0.75,1.4) (see Figure 11). As in the beginning of Section 3, we choose values for the weights *g*_*WT*_ = 0.1 and *g*_*TW*_ = 0.05 using the model output of the average firing rates of the four neuron populations to the model input (shown in Figure 3) as our point of comparison (see Figure 12).

The modified model including connections to and from the mid-brain can capture the delayed inhibition response in the projection neurons from delayed stimulation of the A*β* fibers (see Figure 13).

As concerns neuropathy, we use a similar approach as in Section 3.3 and consider both (a) an inhibitory effect of the A fibers to the C fibers represented by amplitude modulation of the C fibers by the A*β* fibers as given in Equation (5), and (b) circadian rhythm in the C and A*β* fibers. In addition, we use the amended model to investigate the hypothesis that under neuropathy, time spent awake causes increased excitatory input from the mid-brain to the dorsal horn circuit [14]. Thus, we assume that the strength of the connection from the mid-brain population *T* to the projection neuron population *W* given by the weight *g*_TW_ (see Figure 10) increases during wake and decreases during sleep. Moreover, we assume that under normal conditions, the *T* population inhibits the activity of the *W* population, while under neuropathy, the connection from *T* to *W* is excitatory (see red and blue lines in Figure 10). Thus, under normal conditions, the weight *g*_*TW*_ has a daily rhythm shown in red, and under neuropathic conditions, it has the rhythm shown in blue in Figure 14, where negative values result in excitatory input from *T* to *W* because of our choice of using a negative sign in front of *g*_*TW*_ in Equation (6).

**Fig. 11.**
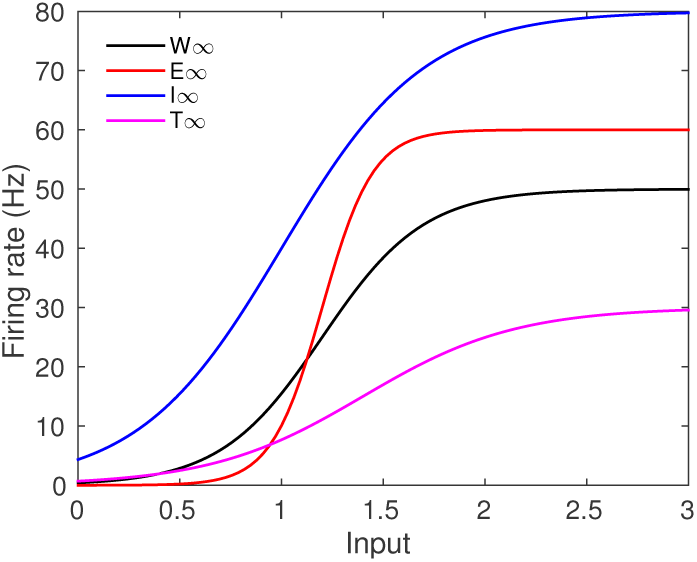
Response functions of the projection (black), excitatory (red), inhibitory (blue), and midbrain (magenta) neuron populations for different constant inputs (on the x-axis).

**Fig. 12.**
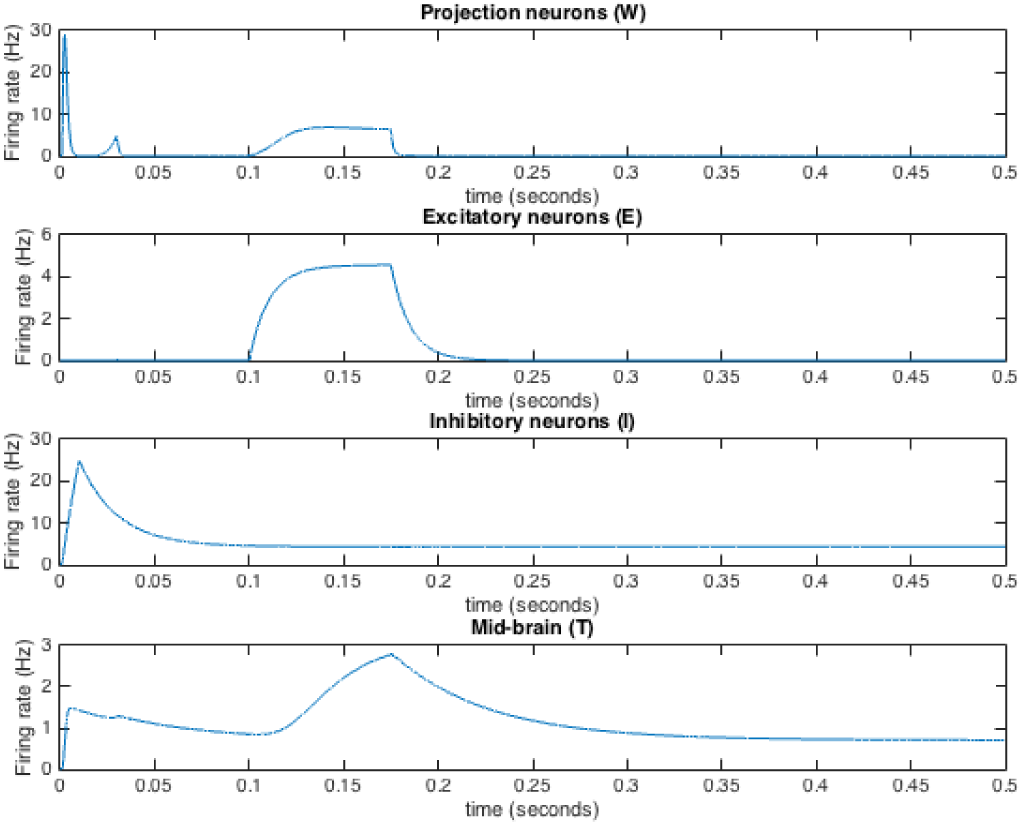
Firing rates for each population (connected as described in Figure 10) in response to the input from afferent fibers as described in Figure 3 including the amendments to the dorsal horn circuit model given in Equation (6).

**Fig. 13.**
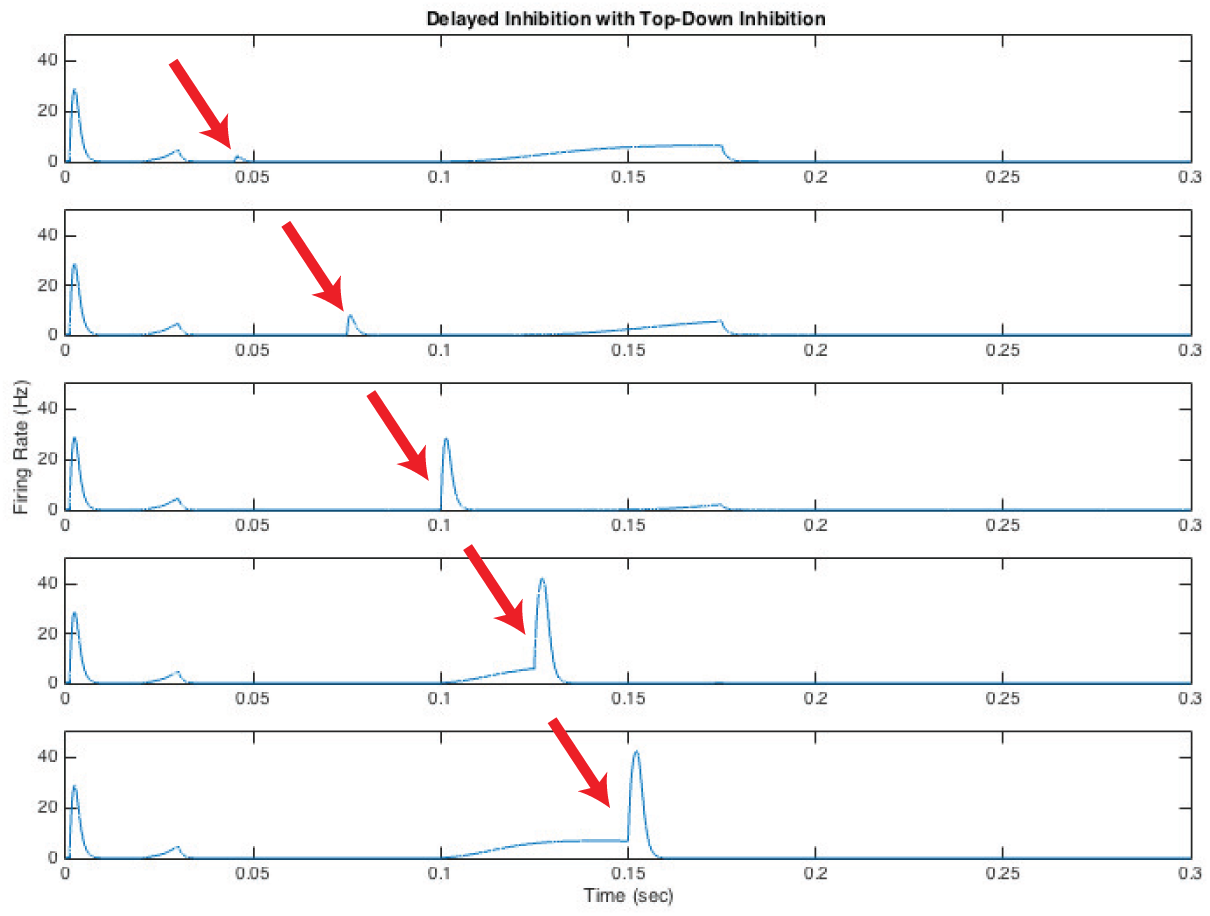
Pain inhibition phenomenon captured in the modified model including top-down inhibition. Response of the projection neuron population to the initial fiber pulse stimulation (at *t* = 0) and the second pulse stimulation only to A*β* fibers (red arrows) for increasing in time between fiber stimulations.

With these above-mentioned modifications to the model of the dorsal horn circuit, our amended model reproduces a more pronounced flip in the phase of the *W* population (see Figure 15) than in the case of only amplitude modulation of the C fibers through A*β* fibers (see Figure 9). We note as well that the W population does not saturate to its maximum firing rate in Figure 15 (right), as it did in the earlier model in Figure 9.

**Fig. 14.**
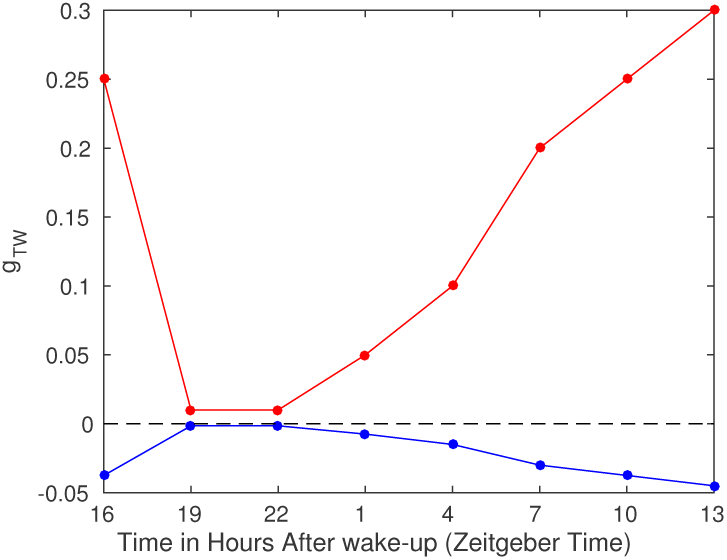
The strength of the connection from the mid-brain to the projection neuron population as a function of hours after wake-up under (blue) neuropathic and (red) normal conditions. We note that negative values of *g*_*TW*_ result in excitatory input from *T* to *W* population, see Equation (6).

## 5 Conclusions and future work

We have constructed a biophysical model of the pain processing circuit in the dorsal horn that represents the interactions between inhibitory and excitatory interneurons, and WDR projection neurons. Our model considers the average firing rate of each of these three neuron populations, and therefore includes less biophysical detail than previous circuit models consisting of single spiking neurons. However, our choice of modeling framework is motivated by our ongoing work to incorporate this model for the pain processing circuit with sleep-wake regulatory network models (see [4] for a review). Such an extended sleep-wake-pain model would allow us to test several existing hypotheses on the effects of sleep-dependent and circadian modulation of pain sensitivity. In addition, we have chosen to use a simplified modeling approach, because it allows us to examine whether suggested mechanisms (i.e., rhythmicity in afferent fibers and their interaction changing from inhibitory to excitatory under neuropathic conditions) are capable of explaining observed rhythmicity in pain before we incorporate more physiological details into our model.

Concerning the phenomenon of wind-up, we simulate it as an increase in the synaptic efficacy of the C fibers before their input reaches the deep dorsal horn circuit. This is an assumption that is supported by experimental evidence of wind-up in the potential of the C fibers observed in the superficial but not in the deep dorsal horn [29]. However, experimental data also suggest that wind-up is more pronounced in the deep than in the superficial dorsal horn neurons [30]. Therefore, our model assumption of wind-up occurring only upstream from the deep dorsal horn is not widely supported by the data. Moreover, at the current stage, our model incorporates no information on possible mechanistic explanations of wind-up. This is a limitation of the model as it restricts our ability to test existing hypotheses (see Figure 6 in [11]) and increase the current understanding of the generation of wind-up.

**Fig. 15.**
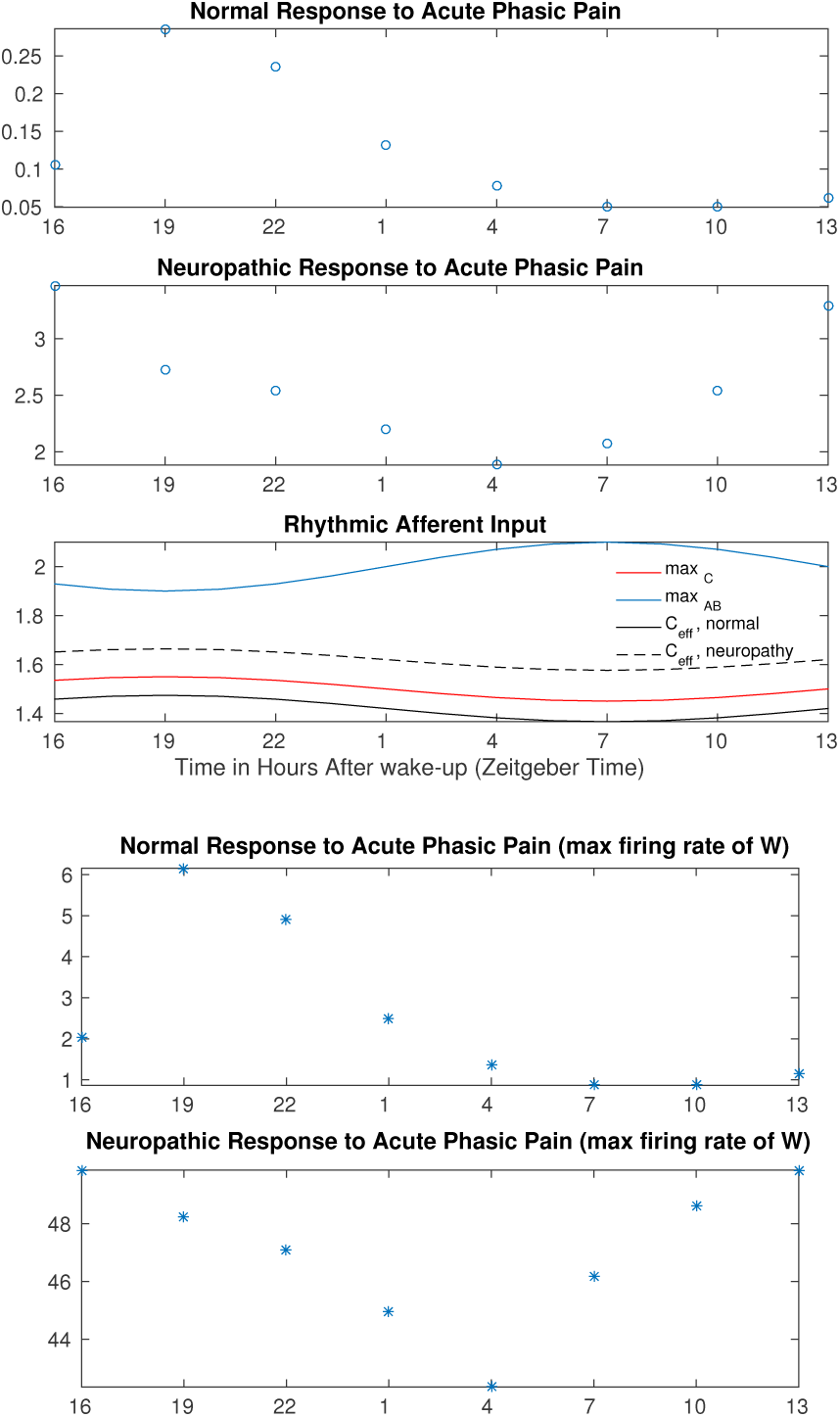
Integrated (circles, top 2 panels) and maximum (asterisks, bottom 2 panels) projection neuron firing rates predicted by the amended equations of time evolution given in (6) under normal (top and 4th panels) and neuropathic (2nd and bottom panels) conditions showing predicted circadian modulation over 24 h. Circadian rhythmicity of responses is generated by the different circadian rhythms in the amplitudes of the afferent fiber-model inputs (middle panel). Under normal conditions, the interneuron population *I* decreases the amplitude of the model input from C fibers (red curve) leading to a reduced effective amplitude of C fiber input (black solid curve). Under neuropathic conditions, the connection between *I* and C fiber model input is excitatory resulting in a higher effective amplitude of the C fiber model input (black dashed curve). We calculate the activity of the projection neuron population as the area under the C-response in the same way as in Figure 9.

As concerns neuropathy, by taking into account both amplitude modulation of the C fibers by the A fibers and normal inhibitory effect switching to excitatory under neuropathic conditions, our model reproduces a change in the daily rhythm seen in the activity of the WDR projection neurons and predicts a higher baseline of pain under neuropathy than under normal conditions, both of which agree with experimental evidence. In the case where a connection to, and from, the mid-brain is included in the dorsal horn circuitry, the flip of the rhythm in the projection neurons is more pronounced and does not evoke a response in the WDR neurons that is at its maximum as in the case where there is no connection to the mid-brain. In our ongoing work, we investigate the robustness of these above-mentioned model predictions for neuropathy, in particular as concerns the range of parameter values that represent the strength of the connection between the WDR projection neurons and the mid-brain.

We have incorporated neuropathy in an attempt to validate that our model can replicate known circadian pain effects. It is important to note that a more biologically realistic model has been developed including large networks of individual neurons [2]. Similarities between our model and the one proposed by Arle et al. is the lack of connection from the A*β* fibers to the inhibitory interneurons, but a major difference is that their model has two distinct circuits for nociceptive and neuropathic pain. We instead use the same circuit but propose different mechanisms within the circuit that contribute to neuropathic pain (for example inhibition switching to excitation under neuropathic conditions). We justify our use of a simplified model by emphasizing that our motivation is in understanding the effect of circadian and sleep-dependent processes on pain sensitivity, and note that our model does capture circadian effects in neuropathic pain patients.

In our ongoing work, we are constructing equations for the time evolution of the average activity of each of the three afferent fiber populations. Such a generalization would not only increase our ability to include possible mechanisms of wind-up but also help in connecting models of pain and sleep together. That is, in the future generalized model, a change in the homeostatic sleep drive (that is an output of the sleep-wake model) could be directly fed into the pain circuit model by influencing the sensitivity of the afferent fibers to external stimuli. This will allow us to more thoroughly investigate several hypotheses on the coupling between sleep deprivation and pain sensitivity.

## Acknowledgements

This work was conducted as a part of A Research Collaboration Workshop for Women in Mathematical Biology at the National Institute for Mathematical and Biological Synthesis, sponsored by the National Science Foundation through NSF Award DBI-1300426, with additional support from The University of Tennessee, Knoxville. This work was additionally partially supported by the following sources: NSF Award DMS-1412119 (VB) and the Pritzker Neuropsychiatric Disorders Research Consortium (MH). Any opinions, findings, and conclusions or recommendations expressed in this material are those of the authors and do not necessarily reflect the views of the National Science Foundation.

